# A Biosensor for Detection of Indole Metabolites

**DOI:** 10.1101/2021.03.05.434140

**Authors:** Jiefei Wang, Chao Zhang, W. Seth Childers

## Abstract

The lack of diverse health-related biosensors limits the progress towards our understanding of how the microbiome metabolism impacts health. Microbially produced indole-3-aldehyde (I3A) has been associated with reducing inflammation in diseases such as ulcerative colitis by stimulating the aryl hydrocarbon receptor (AhR) pathway. We mined the protein database for gut microbiome metabolites’ sensors and developed a biosensor for I3A. We engineered *E. coli* embedded with a single plasmid carrying a chimeric two-component system that detects I3A. Our I3A receptor characterization identified residues that contribute to the sensor’s high specificity in a range of 0.1-10 µM. The I3A biosensor opens the door to sensing indole metabolites produced at the host-microbe interface and provides new parts for synthetic biology applications.

## INTRODUCTION

Synthetic biology has demonstrated the potential for probiotics to address a range of health issues, including the treatment of rare metabolic disorders^*1*^ and pathogenic infections^*2, 3*^. The lack of diverse parts to sense health-related signals limits these applications. We mined the protein database for sensors that could detect signals associated with gut-microbiome interactions.

Amongst microbiome-produced metabolites, indole and its derivatives serve as important interkingdom signaling molecules^*4*^. In the gastrointestinal tract, tryptophan is metabolized by three major metabolic pathways^*5*^: monoamine neurotransmitter serotonin (5-hydroxytryptamine, 5-HT) production^*6*^, nicotinic acid production through kynurenine (Kyn) pathway^*7*^, and production of indole compounds triggering aryl hydrocarbon receptor (AhR) pathway^*8*^. *Lactobacillus*-generated indole-3-aldehyde (I3A), which is detected in gastric fluids and urine, stimulates AhR mediated transcription^*8*^. I3A stimulation of the AhR receptor may have broad health implications that include regulating gut barrier function^*9*^, reprogramming T cells^*10*^, reducing skin inflammation^*11*^, central nervous system (CNS) inflammation^*12*^, and inflammation associated with aging^*13*^. Similarly, pathogenic *E. coli* strains (EPEC, EHEC, and EAEC) secrete certain indole derivatives, including I3A and indole-3-acetic acid (I3AA), to kill *C. elegans*^*14*^. Therefore, I3A is a critical microbially produced biomarker that directly impacts health through AhR activation.

Compared to traditional detection methods, such as mass spectroscopy, cell-based biosensors take advantage of the natural ability of a cell to sense and respond to the environment. Thus cell-based biosensors provide the potential for longitudinal measurements and noninvasive *in vivo* clinical applications^*2, 15, 16*^. Because indole compounds are mainly produced through the metabolism of tryptophan by microbes, we seek to build a sensor for indole compounds in bacteria. While indole derivatives act on eukaryotic AhR^*8, 17*^, those interactions are non-specific and function as a eukaryotic transcription factor, presenting challenges to repurpose AhR as an indole bacterial biosensor. Though indole acts upon bacterial sensor BaeS, CpxA, and EvgS^*18, 19*^, it remains unclear if those indole-receptor interactions are direct or indirect through indole’s impact upon membrane potenial^*20, 21*^. Therefore, the lack of biochemically characterized indole metabolite sensors prevents the use of this biomarker in health applications.

One of the primary biochemical systems microbes use to sense their surroundings is two-component systems (TCSs). TCSs provide a cornerstone of bacterial signaling and coordinate processes such as the cell cycle^*22*^, chemotaxis^*23*^, cell-cell communication^*24*^, and bacterial virulence^*25*^. A TCS consists of a histidine kinase (HK) and a response regulator (RR), which are responsible for sensing environmental signals and regulating cellular processes, respectively (Figure 1A). TCSs are promising targets for engineering biosensors for four reasons^*26*^.

**Figure 1.**
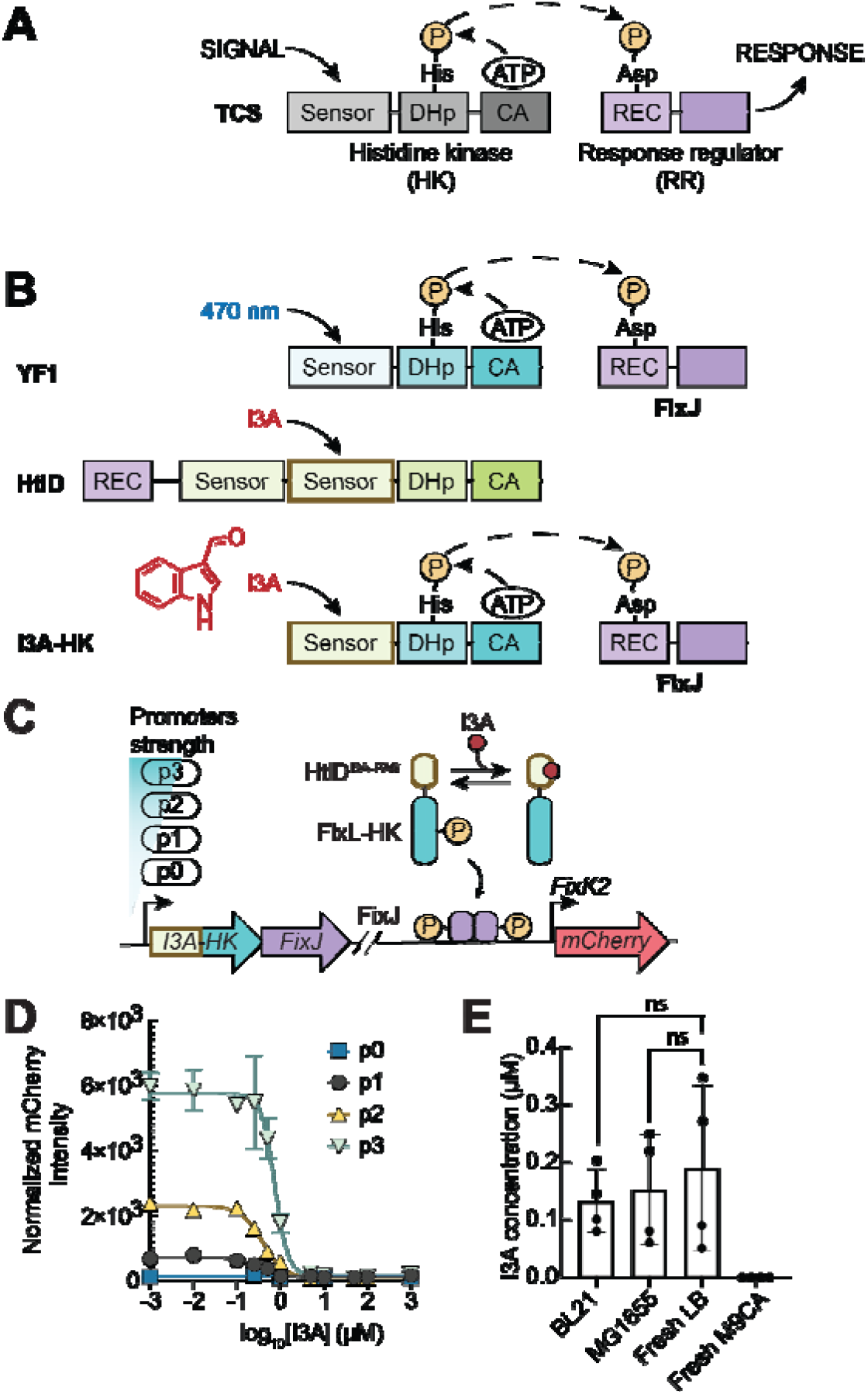
Design of an I3A biosensor. (A) A diagram for a typical two-component system (TCSs). A TCS is composed of a sensor histidine kinase (HK) (grey) and a response regulator (RR) (purple). A conserved HK contains a dimerization histidine phosphotransfer (DHp) domain and a C-terminal ATP-binding catalytic domain (CA) domain. The RR includes a receiver (REC) domain and a DNA binding domain. The signal regulates autophosphorylation of a conserved histidine in the DHp domain. Further phosphotransfer from the DHp domain to the REC domain results in downstream gene regulation. (B) Design of an I3A sensing chimeric histidine kinase I3A-HK. YF1 is a chimeric protein that consists of a light-sensing PAS domain and FixL-HK domain, the conserved kinase core from FixL^*56*^. The HtlD^I3A-PAS^ domain, an I3A binding PAS domain, is the second PAS domain in the HtlD as indicated (brown box). The chemical structure of I3A is shown (red). We swapped the light-sensing domain in YF1 to the HtlD^I3A-PAS^ and generated the chimeric protein I3A-HK. (C) The construct pDusk-I3A_p0-p3 for I3A-repressed *mCherry* expression. I3A regulates the activity of I3A-HK in phosphorylating the FixJ RR, which stimulates transcription of *mCherry* at the *FixK2* promoter. I3A repressed expression of *mCherry*. (D) I3A dose-response curve for pDusk-I3A_p0-p3. Normalized fluorescent intensity to OD_600_ was plotted against the concentration of I3A. See the curve fitting results in Table 1 for the three biological replicates. (E) The quantification of I3A extracted from culture media and saturated *E. coli* strains as determined by HPLC-MS. The I3A concentrations are plotted in the bar chart. Data represent the mean ± SD of at least 3 biological replicates. ns: not significant.

First, TCSs provide a warehouse of sensors^*27*^, including pH, metals, quorum sensing compounds, and secondary metabolites. For instance, TCSs sensing a broad spectrum of light are used as optogenetic tools in *E*.*coli*^*28-32*^. TCS sensing thiosulfate has been developed and applied in *E*.*coli* for diagnosing colon inflammation in mice^*3, 33*^. Also, TCS metabolic sensing has been developed to screen unique chassis producing small molecules and feedstock^*33-37*^. Despite significant HK characterization progress, the sensing signals remain unknown for most HKs^*38-40*^. The fact that signals remain unknown is partially due to the lack of crystal structures of sensors co-crystallized with their stimulating ligands.

Second, TCSs have remarkable plasticity in how signals flow within and between the HK and RR. On the one hand, the sensing domain can be swapped with homologous sensory domains to sense different signals^*41, 42*^. On the other hand, TCSs also exert diverse outputs that include regulation of transcription, translation, motility, and proteolysis^*43, 44,45*^. Additionally, the HK-RR protein-protein interaction interface can be engineered to alter phosphate transfer to other response regulators^*46*^. Therefore, the two-component systems’ modularity allows for the straightforward and customized rewiring of a signal input to the desired signaling outputs.

Third, productive cross-talk amongst several TCSs can function as an integrated multi-kinase network (MKN)^*47*^. MKNs allow sensing multiple signals simultaneously and minimize the chance of off-target regulation in a physiological environment. For example, to improve the stringency of uranium sensing, two functionally independent TCSs, UzcRS, and UrpRS, were assembled as a AND gate^*48*^.

Fourth, TCSs are transferable to eukaryotes^*49, 50*^, which expand engineered biosensors’ potential to be applied in prokaryotic and eukaryotic cells. Here we identified an I3A binding domain and engineered a chimeric HK that regulates mCherry expression upon the addition of I3A in *E. coli*. We engineered two one-plasmid systems, which carry the I3A binding domain within a chimeric histidine kinase and its cognate response regulator. This initial indole metabolite sensor provides a scaffold in synthetic biology for engineering sensors for a diverse repertoire of indole metabolites. This includes metabolites such as indole, which is known to induce virulence factors, promote resistance to pathogens, and promote homeostasis of gut epithelial function^*5, 8, 51*^.

## RESULTS & DISCUSSION

### Identification of an indole-3-aldehyde binding PAS domain

We mined the protein database (PDB) for bacterial sensory domains that interact with signals in human microbiome environments. We focused our search upon Per-Arnt-Sim (PAS) domains, as they are essential sensing modules that commonly regulate histidine kinases. We found 1293 PAS domain entries and identified 38 entries that bind unique ligands (Table S1).

Amongst ligand-associated PAS domains, we identified a PAS domain co-crystallized with I3A (PDB: 3BWL)^*52*^. I3A plays a vital role in maintaining mucosal homeostasis by stimulating the AhR signaling pathway^*5, 8*^. The I3A-binding PAS domain belongs to the protein HtlD (halobacterial transducer of rhodopsin (HTR)-like protein, UniProt Q5V5P7)^*53*^ encoded by *rrnAC0075* in halophilic Archaeon *Haloarcula marismortui*. Accordingly, we named the I3A binding PAS domain as HtlD^I3A-PAS^ in this work and characterized the ability of HtlD^I3A-PAS^ to serve in an I3A biosensor.

### Engineer a two-component system in E.coli to sense indole-3-aldehyde

The HtlD signaling protein consists of a receiver domain (REC) followed by an HK domain (Figure 1B). However, there are no cognate RRs or histidine phosphotransfer proteins (Hpt) within the proximal genomic regions. The lack of knowledge of the downstream RR and the regulating set of genes prevented our ability to refactor the entire signaling system in *E. coli*. To overcome this problem, chimeric HKs have demonstrated the ability to exchange sensory domains for producing biosensors^*41, 42*^. One such example is the blue light-sensing chimeric histidine kinase YF1^*42, 54, 55*^. The sensory domain of the blue light-sensing YF1 is also a PAS domain (YF1^LOV-PAS^), which is structurally similar to the HtlD^I3A-PAS^ (Figure S1A). Therefore, to exploit the I3A sensing capabilities of HtlD^I3A-PAS^, we swapped the LOV sensory domain from YF1 with the HtlD^I3A-PAS^ and tested if the chimeric HK could sense I3A.

To determine the optimal sensor-HK chimera fusion site, we exploited a set of conserved D-(A/I/V)-(T/S)-E residues at the C-terminus of the PAS domain fold. This conserved motif forms a network of hydrogen bonds with the central beta-sheets of the PAS domain fold^*56, 57*^. As demonstrated with the YF1 chimera construction^*56*^, this C-terminal motif in PAS provides a conserved modular point to swap PAS sensory domains. Therefore, we swapped the YF1^LOV-PAS^ to its DIT motif with the HtlD^I3A-PAS^ up to its DIT motif and named the chimeric protein I3A-HK (Figure 1B, Figure S1B).

We introduced the I3A-HK into two plasmids, pDusk and pDawn^*54*^, which contain well-characterized parts downstream of the sensor and exhibits low background in the absence of blue light^*54*^. We designed and constructed plasmid pDusk-I3A_p1. Within the plasmid pDusk-I3A_p1, a bicistronic operon, regulated by the promoter p1, controls the constitutive expression of the I3A-HK and the paired RR. I3A regulates phosphorylation of the RR, then triggers transcriptional repression of the mCherry reporter gene^*54, 56*^ (Figure 1C). To invert the signal so that the introduction of I3A increases the reporter gene expression, we modified the pDawn plasmid^*54*^ to incorporate the I3A-HK resulting in the plasmid pDawn-I3A (Figure S2A). In the engineered pDawn-I3A, I3A represses the expression of the λ phage repressor cI, which represses the expression of mCherry through the λ promoter *pR*^*54, 58*^.

Upon addition of 10 µM I3A to strains containing the pDusk-I3A_p1 plasmid, we observed no impact upon cell growth (Figure S2B) and a 3-fold decrease in mCherry expression relative to cultures supplemented with DMSO (Figure S2B). An I3A titration yielded a dose-response curve with an IC_50_ of 0.3 ± 0.1 µM (Figure 1D, Table 1). Similarly, upon the addition of 10 µM I3A to strains containing the pDawn-I3A_p1, we observed a 2.5-fold increase in mCherry expression. Further I3A titration studies yielded a dose-response curve with an EC_50_ of 11.5 ± 7.8 µM (Figure S2C, Table 1). Therefore, we have constructed two biosensors that repress or stimulate output gene expression upon the addition of I3A. Notably, the pDawn-I3A system exhibited a higher EC_50_ than the IC_50_ of the pDusk-I3A system (Table 1). We suspect that the difference is caused by the involvement of the λ phage repressor cI in pDawn-I3A. Other than the involvement of cI, the difference may be caused by the dynamic range of the pDusk-I3A_p1. Given that the dynamic range of the pDusk-I3A_p1 is narrow and the fluorescent intensity is low even without I3A, a high concentration of I3A may lead to cut-off distortion in detection.

**Table 1.**
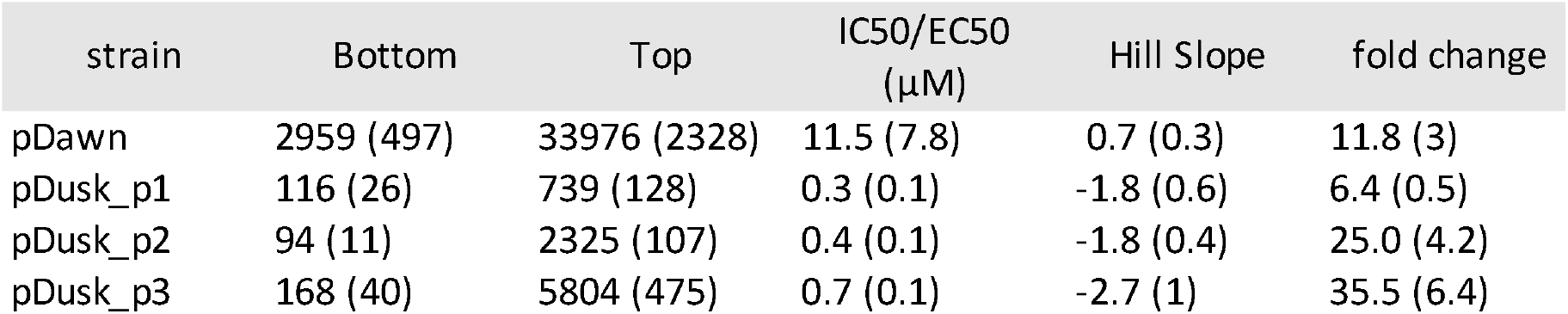
Fitting for the dose-response curves. Each curve is fitted to the Hill equation with default settings in Prism9. Bottom: predicted minimal mCherry; Top: predicted maximal mCherry. Standard deviation estimates are in parentheses. The mean and SD are reported in the table for at least 3 biological replicates.

| strain | Bottom | Top | IC50/EC50<br>(μM) | Hill Slope | fold change |
| --- | --- | --- | --- | --- | --- |
| pDawn | 2959 (497) | 33976 (2328) | 11.5 (7.8) | 0.7 (0.3) | 11.8 (3) |
| pDusk_p1 | 116 (26) | 739 (128) | 0.3 (0.1) | -1.8 (0.6) | 6.4 (0.5) |
| pDusk_p2 | 94 (11) | 2325 (107) | 0.4 (0.1) | -1.8 (0.4) | 25.0 (4.2) |
| pDusk_p3 | 168 (40) | 5804 (475) | 0.7 (0.1) | -2.7 (1) | 35.5 (6.4) |

### Tuning the dynamic range of the I3A sensor

One approach to improving the dynamic range is to tune the ratio of receptor and substrate concentration^*31*^. To adjust the receptor to substrate ratio, we varied the promoter of I3A-HK using a set of well-characterized promoters from the BIOFAB kit^*59*^ (Figure 1C, Table S2): p0, p2, and p3. We grew *E. coli* cultures carrying constructed plasmids in the presence of DMSO or I3A at various concentrations to determine the effect of I3A upon the mCherry expression (Figure 1D). The dose-response curves were fitted to the Hill equation, as shown in Table 1. Replacing the promoter enlarged the dynamic range, consistent with the strength of the promoter region (Table S2), with observed fold changes of 6.4-fold for p1, 25-fold for p2, and 35.5-fold for p3. The IC_50_ is 0.4 ± 0.1 µM for p2 and 0.7 ± 0.1 µM for p3, similar to the IC_50_ for p1. The weakest selected promoter (p0) displayed no observable changes in mCherry expression upon the addition of I3A. Therefore, we demonstrated that the response to I3A could be tuned predictably by varying the promoter strength for the I3A-HK. Our ability to accurately sense I3A could be affected by endogenous production of I3A by *E. coli* or the abundance of I3A in LB media. Indeed, co-crystallization of the sensor-I3A complex indicates that I3A likely interacts with HtlD^I3A-PAS^ during protein expression^*52*^. Moreover, past experiments suggest that *E. coli* might produce I3A^*14*^. To interrogate how much I3A is in the growth culture, we measured the I3A concentration in the cell culture media via HPLC-MS. We observed that freshly prepared LB media contained 0.19 ± 0.07 μM of I3A, while minimal M9 media contained no detectable amounts of I3A.

We recovered 0.13 ± 0.03 μM of I3A from *E*.*coli* BL21 and 0.15 ± 0.05 μM of I3A from *E*.*coli* MG1655 wild-type strains. The I3A concentration from both BL21 and MG1655 wild-type strains are similar to the amounts of I3A in fresh LB media, 0.19 ± 0.07 μM (Figure 1E, Figure S2D-F). Therefore, we observed no additional I3A production by *E* coli under the tested growth conditions. However, it is possible that I3A production could be condition or strain-dependent^*14*^. Therefore, the endogenous I3A in fresh LB media provides a minor 0.19 ± 0.07 μM offset to our calculated IC_50_ and EC_50_. The unexpected observation of I3A in the LB media (Figure 1E) may warrant its future investigation of I3A as a potential yeast secondary metabolite or the source of its contamination.

### Interrogation of the I3A receptor binding site

A key question in part characterization is if structural features of the I3A chimeric sensor function as expected. PAS domains contain a conserved hinge motif D-(A/I/V)-(T/S) that couples the conformational change within the PAS to the conformational re-arrangement that allows HK autophosphorylation^*60-62*^. Therefore, we mutated the DIT motif of the I3A-HK gene to AIA (i.e., D507A and T509A) to interrogate if this hinge motif was also critical for I3A sensing. We observed that the DIT to AIA mutation in the pDusk-I3A_p1 plasmid diminished the responsiveness to I3A up to 10 µM (Figure 2A, Figure S2G). This indicates that the DIT motif mutants disrupt signaling, which may be due to a lack of allostery between the sensor and HK as observed in other HKs^*56, 63, 64*^. However, we cannot rule out that the double mutant caused protein misfolding. The data indicate that the I3A stimulation of mCherry expression requires a functional PAS sensory domain.

**Figure 2.**
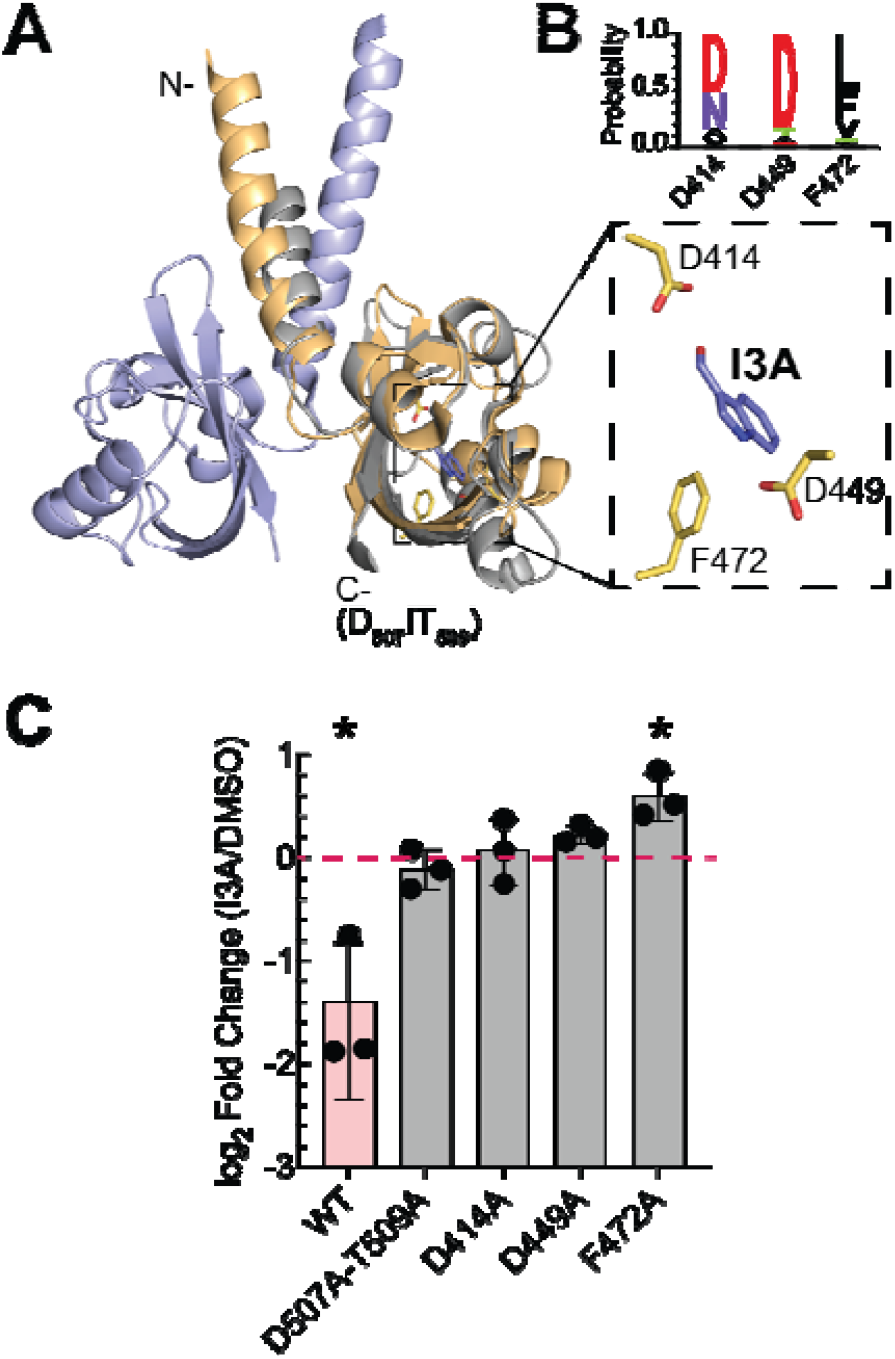
The I3A sensing through PAS domain. (A) Structure of the I3A binding PAS domain, HtlD^I3A-PAS^ (PDB: 3BWL) deposited by Osipiuk, J., Zhou, M., Freeman, L., and Joachimiak A. at the Midwest Center for Structural Genomics^*52*^. Dimer of HtlD^I3A-PAS^ is shown (light orange and light blue). The structure of HtlD^I3A-PAS^ is aligned to the light-sensing PAS domain of YF1 (PDB: 4GCZ) (grey). The signal transmission residues are located at C-terminal in the presented structure. The residues within 5 Å to the I3A are indicated (dashed box). (B) Conservation of the potential I3A binding residues are determined by sequence alignment and plotted using WebLogo^*74, 75*^ as described in the method. (C) I3A binding site (D414A, D449A. F472A) and allosteric signal transmission residues (D-(A/I/V)-(T/S)-E as D507A-T509A) were mutated as indicated on the pDusk-I3A_p1 plasmid. Fold changes are plotted in log scale. Fold change is measured as the ratio of normalized mCherry intensity of I3A-treated vs. DMSO-treated. Ratio equals to 1 is labeled (red dash line). Data represent the mean ± SD of at least 3 biological replicates. *: p < 0.05, comparing I3A-treated vs. DMSO-treated samples.

We next tested how changes to the I3A PAS domain binding site residues impacted I3A sensing. We examined three interacting residues within 5 Å of the I3A ligand: D414, D449, and F472 (PDB: 3BWL)^*52*^ (Figure 2A-B). We mutated D414, D449, and F472 within the I3A-HK gene individually to alanine in the pDusk-I3A_p1 plasmid using site-directed mutagenesis and interrogated I3A-regulated mCherry expression (Figure 2C, Figure S2G). D414A and D449A I3A-HK variants were not responsive to I3A up to a concentration of 10 µM. In comparison, rather than reduction, the F472A I3A-HK variant resulted in 1.5-fold induction of mCherry. This observed inversion of the response to I3A by the F472A variant, suggests that F472 may be involved in allosterically coupling ligand binding to changes in the HK conformational state. Overall, the interrogation of the I3A receptor binding site confirmed that D414, D449, and F472 are critical for I3A sensing. Future engineering of the amino acid positions present opportunities to improve I3A-binding affinity for the sensor or alter the substrate specificity to other closely related indole metabolites.

Interestingly, in the presence of DMSO, the I3A binding site mutants (D414, D449, and F472) exhibited a 3 to 10-fold increase in mCherry expression relative to the wild-type I3A-HK (Figure S2G). This increased mCherry expression could be due to increased expression levels of the I3A-HK binding site mutants, similar to our observation of promoter tuning (Figure 1). Although, each I3A-HK variant used the same promoter. Alternatively, the observed I3A in the LB media by HPLC-MS (Figure 1E) may repress mCherry expression in the wild-type I3A-HK sensor but may be unable to stimulate the I3A-HK binding site mutants (Figure S2G). We envision the I3A biosensor will provide a powerful tool to identify the bacterial I3A biosynthetic pathway in bacteria suspected to produce I3A^*8*^.

### Selective I3A sensing through PAS domain

To examine the specificity of the I3A sensing, we treated the engineered *E. coli* against a panel of 11 closely related indole metabolites at a concentration of 10 µM and 100 µM. Many of these indole metabolites only differ by one functional group where the R_1_-R_4_ group is substituted from -CHO in I3A to other groups (Figure 3A). Amongst this set, only I3A could stimulate mCherry expression at a concentration of 10 µM (Figure 3B). We observed a similar specific I3A stimulation of mCherry expression from the pDawn-I3A plasmid (Figure S3A) as from pDusk-I3A-p3 plasmid (Figure S3B). Lack of stimulation by the other closely related indole metabolites could be due to a lack of binding to the I3A sensor or the inability of the indole metabolites to cross the cell wall. At a 100 µM ligand concentration, we observed a mild but statistically significant reduction of mCherry expression of less than 10% by the following: I3M, I3AA, IND, I7OH, and IMI (Figure S3B). These results indicate that I3A is a potent and specific stimulator of I3A-HK. While I3M, I3AA, IND, I7OH, and IMI weakly stimulate mCherry expression at concentrations 100-fold greater than I3A mediated stimulation of I3A-HK.

**Figure 3.**
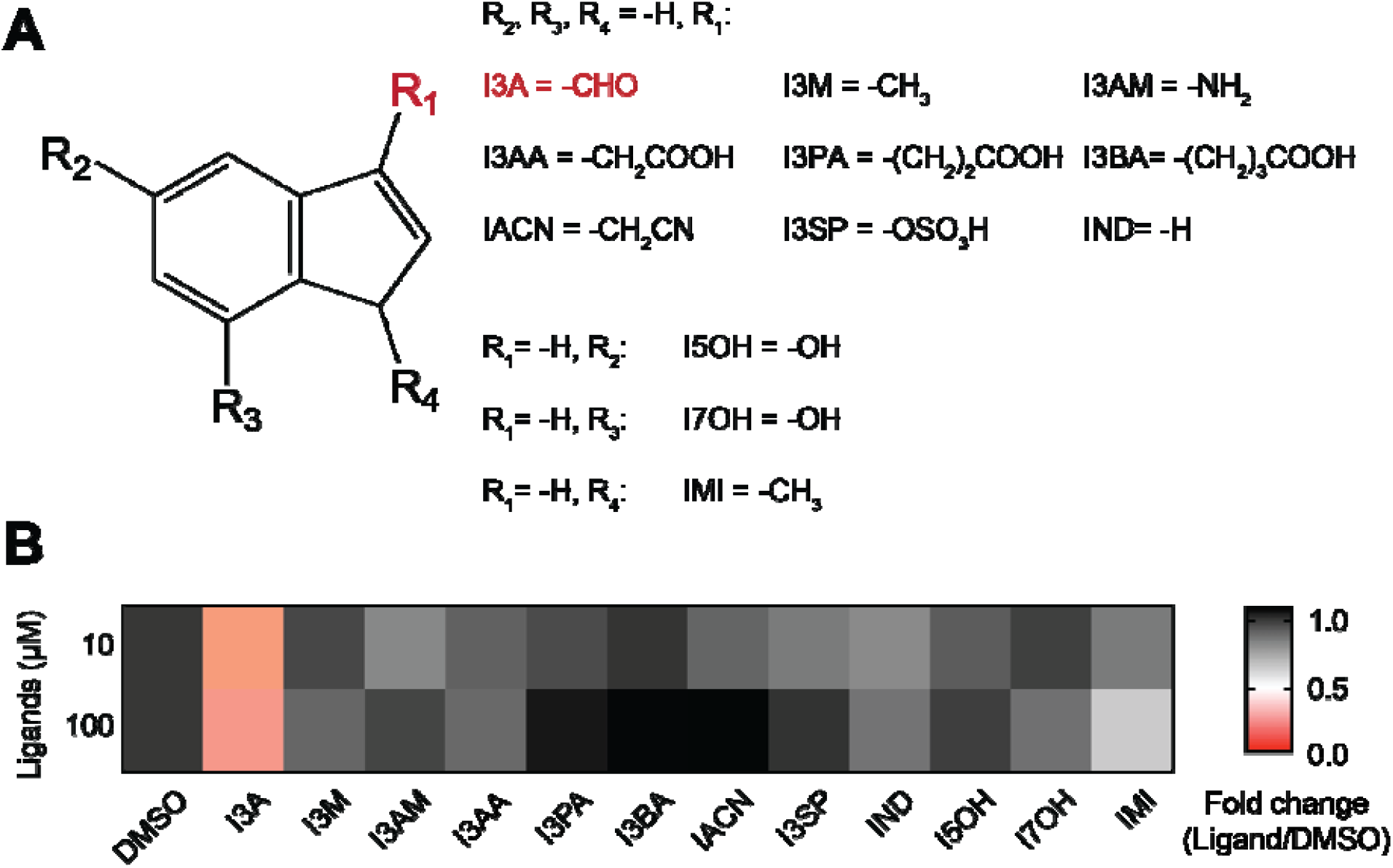
The selectivity of the I3A sensor. (A) Structure of indole derivatives that were used in the selectivity tests. I3A: indole-3-aldehyde, I3M: 3-methylindole, I3AM: 3-aminoindole, I3AA: Indole-3-acetic acid, I3PA: indole-3-pyruvic acid, I3BA: Indole-3-butyric acid, IACN: 3-Indoleacetonitrile, I3SP: indoxyl sulfate, IND: indole, I5OH: 5-Hydroxyindole, I7OH: 7-Hydroxyindole, IMI: 1-Methylindole (B) The selectivity of pDusk-I3A_p3. The fold-change is reported as a ratio of normalized mCherry intensity for ligand-treated vs. DMSO-treated samples. The fold-change is plotted as a heat map given the gradient as shown in the figure, with a max (1.1 fold, black) and a min (0 fold, red). Data represent the mean of at least 3 biological replicates. DMSO: dimethyl sulfoxide. See the corresponding bar charts and the results of the test for the pDawn-I3A in Figure S3A.

Notably, indole has been the most studied amongst these metabolites and reported at a much higher stimulating concentration^*20*^. We measured indole concentration via LC-MS and found *E. coli* BL21 strain produce 237.5 ± 1.1 µM indole in LB media (Figure S3C-D). However, the amount of indole endogenously produced by the *E. coli* BL21 strain expressing our chimeric I3A-HK did not stimulate complete repression of mCherry expression. Therefore, we tested higher exogenous addition of indole concentrations in the range of 500-1000 µM and observed repression of mCherry expression (Figure S3E). These data suggest that the I3A-HK has high specificity for I3A and a 100-1000-fold higher IC_50_ for indole. The indole detection could be through direct binding with the I3A sensor, or exogenous indole could stimulate biosynthesis of I3A in *E*.*coli*^*14*^. However, we also suspect that high indole concentrations used in the indole titration experiment could induce mild toxicity (Figure S3G) that impacts translation. Further genetic and biochemical studies will be needed to distinguish between direct and indirect indole stimulation models.

In addition, we observed that 100-1000 µM I3AA could be detected by the I3A-HK sensor (Figure S3F and H). The weak indole and I3AA repression indicate that one should cautiously consider conditions known to produce high levels of indole or I3AA^*20*^, as it may provide a false-positive result for I3A detection. Critically, the mild sensing promiscuity suggests that enzyme engineering approaches upon the HtlD^I3A-PAS^-like PAS domains offer the potential to alter its specificity for I3A towards other indole metabolites such as IND or I3AA.

### Homologs of the indole-3-aldehyde binding PAS domain

It is not clear why *Haloarcula marismortui* senses I3A and how the regulation through I3A benefits this organism. HtlD^I3A-PAS^ homologs exist in archaea and bacteria (Figure S4A and Table S3), and we investigated if I3A sensing was conserved amongst the HtlD^I3A-PAS^ homologs PAS-1 to PAS-3 (Figure 4 and Figure 4B-D). We found each homologs’ specificity corresponds to their sequence similarity to HtlD^I3A-PAS^. We observed that PAS-2-HK could repress the expression of mCherry upon the addition of I3A with an IC_50_ of 1.1 ± 0.4 µM (Figure S4E). Similarly, expression of mCherry could be weakly repressed by the PAS-3-HK upon the addition of I3A with an IC_50_ of about 0.2 ± 0.1 µM (Figure S4E). Notably, both PAS-2-HK and PAS-3-HK exhibited a smaller dynamic range than their parent strain pDusk-I3A_p3 (Figure S4E). Critically, our validation of a direct I3A binding PAS domain that is conserved in a subset of archaea raises questions about the potential role of I3A signaling in archaea^*65*^.

**Figure 4.**
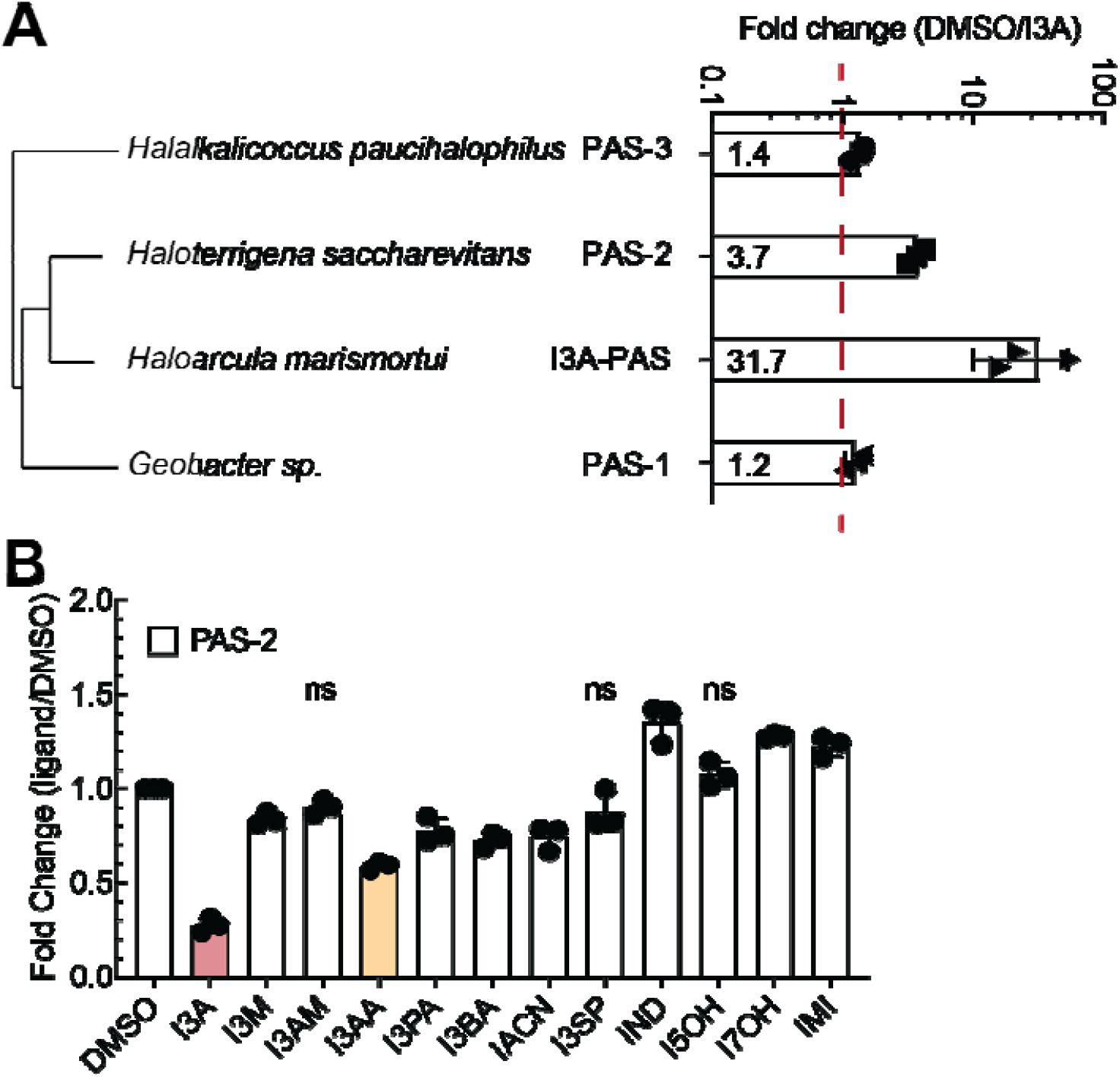
I3A detection is conserved amongst homologs of HtlD^I3A-PAS^. (A) Response to I3A of the homologs of HtlD^I3A-PAS^. To test the homologs, we swapped the HtlD^I3A-PAS^ in pDusk-I3A_p3 to each homolog and generated constructs for chimeric PAS-HKs. See the sequences of homologs in Table S3. The fold-change is reported as a ratio of normalized mCherry intensity for DMSO-treated vs. I3A-treated samples. Data represent the mean ± SD of at least 3 biological replicates. The mean is indicated in a column for each homolog. See a full phylogenetic tree in Figure S4A. (B) The selectivity of PAS-2. The fold-change is reported as a ratio of normalized mCherry intensity for ligand-treated vs. DMSO-treated samples. Data represent the mean ± SD of at least 3 biological replicates. The conditions where ligand-treated vs. DMSO-treated gives a p > 0.05 are indicated with ns. The ligands that give a >1.5-fold change in mCherry expression are I3A (red) and I3AA (yellow). DMSO: dimethyl sulfoxide. See the results for the selectivity of PAS-1 and PAS-3 in Figure S4.

In summary, we have developed a biosensor to detect I3A with high specificity, demonstrating the PDB provides a ripe untapped source for biosensor design. We also identified critical residues surrounding the I3A binding pocket, which will provide the foundation for engineering changes in specificity to detect other health-associated biomarkers such as indole and I3AA. We envision this new I3A biosensor will open the door to new cell-based approaches to interrogate the indole signaling landscape in diverse microbiome environments. Moreover, the I3A biosensor could be a critical new part for cell-based therapeutics^*66*^ that could respond to I3A or even alter the indole signaling landscape^*67*^.

## MATERIAL AND METHODS

### Chemicals

Indole and its derivatives (Table S8) used in this work were purchased from ACROS organics™ and TCI America™. Chemicals were stored in appropriate conditions after opening. DMSO was used to prepare a 100 mM stock solution for each ligand and frozen in a −20 °C freezer until ready to use. In the assay for measuring dynamic range, 100 mM stock solution was diluted into a few lower concentration stocks in DMSO before adding to bacterial liquid culture.

### Bacterial strains and growth

All *E*.*coli* strains, plasmids, plasmid construction methods, primers, and enzymes used in this study are listed in Table S4-6. Primers used for the construction of plasmids were designed using j5 software^*68*^. All primers were purchased from Integrated DNA Technologies (IDT, Coralville, IA). All PAS domains were synthesized as gBlocks from IDT and replaced the YF1^LOV-PAS^ within the pDusk plasmid. The sequence for the chimeric site is shown in Figure S1B. For testing promoters, pDusk-I3A was used as templates and treated as p1. The 200 bp region upstream the I3A-HK in pDusk-I3A was replaced with the selected promoter and ribosomal binding site (RBS) regions. The selected regions were cloned from BIOFAB kit strains B7, B11, and B12 for p0, p2, and p3, respectively (Drew Endy Lab, Stanford University, USA)^*59*^. See Table S2 for sequences. All constructs above were assembled via Gibson Assembly using standard protocols^*69*^. The plasmids pDusk-I3A and pDusk-I3A_p3 were used as templates to generate single-site mutations. Mutagenesis was done with QuickChange. Plasmids were routinely isolated using the GeneJET Plasmid Miniprep kit (Thermofisher, USA).

*E*.*coli* strains (Table S6) were grown in Luria-Bertani (LB) media or on LB plates, including 1.5% w/v agar. Antibiotics were included at the following final concentrations: ampicillin, 100 mg/L, kanamycin, 30 mg/L or chloramphenicol, 30 mg/L.

### Fluorescence reporter assay

Plasmids were transformed into chemical BL21(DE3) *E. coli* cells. Cells were plated onto selective LB media plates and grown overnight at 37 ° C. From a single colony, 2 mL cultures were initiated and incubated at 37 ° C with shaking at 250 rpm for 12 h. The saturated cultures were normalized to OD_600_ = 1 and were inoculated at 1% v/v to 1 mL selected LB media. The stock solution of ligands prepared as different concentrations in DMSO was added to the cultures at 1% v/v upon inoculation. Cultures were incubated at 37 ° C with shaking at 250 rpm. After 20 h^*54*^, 150 µL from each culture was aliquoted into a black-bottom 96-well plate (Greiner Bio-One, USA) for fluorescence measurements, and 50 µL were aliquoted in 100 µL LB media into a clear 96-well plate for measurements of optical density. All endpoint assays were measured in 96 well plates using the Infinite^®^ M1000 plate reader (Tecan, Switzerland). mCherry fluorescence was measured using an excitation wavelength of 585 nm, an emission wavelength of 610 nm, a fixed gain of 60, and a bandwidth of 5 nm. 0.02mg/mL TAMRA SE dye (ThermoFisher, USA) was diluted and used as a standard instrumental control. The optical density was measured at 600 nm (OD_600_). The ratio of fluorescent intensity to OD_600_ was recorded as normalized mCherry intensity. All the experiments were done with at least three replicates using cultures from different colonies. The mean and standard deviation of the normalized mCherry intensity were reported.

### Fitting to the Hill equation

The measure fluorescent intensity was normalized by optical density for plotting and fitting in Prism9 (GraphPad Software, San Diego, USA). The normalized mCherry intensity at each ligand concentration were fitted to the Hill equation in Prism8 with default parameters, using [inhibitor] vs. response - variable slope function with Y = bottom + (top-bottom)/(1+(IC50/X)^hillslope), or [agonist]] vs. response - variable slope function with Y = bottom + (X^hillslope)*(top-bottom)/(X^hillslope + EC50^hillslope). Bottom and top are minimal and maximal response, IC_50_ or EC_50_ is the concentration of ligands that gives a response halfway between top and bottom, and hillslope describes the steepness of curves.

### Method for indole compounds quantification by LC-MS

Quantification of indole compounds was performed using the modified method previously published^*70*^. Individual *E. coli* colony was picked on an LB agar plate and inoculated into a 5 mL LB culture overnight. To extract, the overnight culture was centrifuged at 4000 g for 10 minutes at 4 °C, and the cell pellet and supernatant were separated. The pellet was washed with 2 mL of MilliQ water 3 times to remove residual compounds. Then 0.5 µL Lysonase Bioprocessing Reagent (MilliPore Sigma) was added to the pellet and incubate at room temperature for 5 minutes. The supernatant was transferred into another 15 mL conical tube, combined with lysed cells, mixed with 5 mL of ethyl acetate (FisherScientific, USA), and mixed vigorously. The mixture was then allowed to stand for the layers to separate. The extraction was repeated twice, and the organic layers from each extraction were combined. The organic layer was dried with anhydrous Na_2_SO_4_ (FisherScientific, USA) and evaporated to dryness using a rotary evaporator. The crude extract was dissolved in 1 mL LC-MS grade methanol (FisherScientific, USA), centrifuged to remove particles, filtered with PTFE membrane (FisherScientific, USA), and analyzed by LC-HRMS using SIM mode on Thermo Scientific Q-Exactive Orbitrap (ThermoFisher, USA). A gradient from 50-80% acetonitrile (FisherScientific, USA) over 30 min was used as a mobile phase using an Acclaim Polar Advantage II C18 column (Thermo Fisher Scientific, USA). For quantification of indole-3-aldehyde and indole, a standard curve that correlates the area with the analyte concentration was used. To measure the extraction efficiency through this method, a known amount of analyte was added to water, followed by the above-described method. The efficiency is expressed as the amount calculated from the LC-MS quantification method over a known amount, which is 96.7 ± 0.7% for I3A and 97.6 ± 0.3% for IND. All biological samples were repeated at least three times as replicates.

### Bioinformatics analysis

The alignment of HtlD^I3A-PAS^ (UniProt: https://www.uniprot.org/uniprot/Q5V5P7)^*53*^ was performed (12/2019) using XtalPred (http://xtalpred.godziklab.org/XtalPred-cgi/xtal.pl)^*71, 72*^. The alignment was queried against the UniProtKB database with default settings (Table S7). The result was downloaded and filtered to include 40% identity and 70% coverage, resulting in 25 proteins. The sequences of the 25 proteins were downloaded from the UniProtKB database. The alignment figure was generated using Jalview (http://www.jalview.org)^*73*^ and WebLogo (http://weblogo.threeplusone.com)^*74, 75*^.

## ABBREVIATIONS

5-HT: 5-hydroxytryptamine
AhR: Aryl hydrocarbon receptor
CA: Catalytic domain
CNS: Central nervous system
DHp: Dimerization histidine phosphotransfer
DMSO: DMSO (Biological Grade)
HK: Histidine kinase
HtlD: HTR-like protein D
HTR: Halobacterial transducer of rhodopsin
I3A: Indole-3-aldehyde
I3AA: Indole-3-acetic acid
I3AM: 3-aminoindole
I3BA: Indole-3-butyric acid
I3M: 3-Methylindole
I3PA: indole-3-pyruvic acid
I3SP: 3-Indoxyl sulfate
I5OH: 5-Hydroxyindole
I7OH: 7-Hydroxyindole
IACN: 3-Indolylacetonitrile
IMI: 1-Methylindole
IND: Indole
Kan: Kanamycin
Kyn: Kynurenine
LB: Luria-Bertani
MKN: Multi-kinase network
PAS: Per-Arnt-Sim
PDB: Protein database
RBS: Ribosomal binding site
REC: Receiver domain
RR: Response regulator

## ACKNOWLEDGEMENT

We thank Osipiuk, J, Zhou M., Freeman L., Joachimiak, A., and the Midwest Center for Structural Genomics for their past work structurally characterizing the I3A sensor (PDB: 3BWL). Their tireless structural characterization efforts have created a valuable resource for the scientific community. We thank the Andreas Möglich lab at the University of Bayreuth and Addgene for providing the pDusk (Addgene Plasmid # 43795) and pDawn plasmid (Addgene Plasmid # 43796) that was used in this study. We thank Drew Endy lab at Stanford University and Addgene for the BIOFAB promoter library (Addgene Kit #1000000037). We thank the Yiming Wang lab at the University of Pittsburgh for providing IMI and indole used in this study and Yidong Wang and Qin Zhu’s generous help with bacterial cell lysate extract. We thank Xiaole Yang of the Childers laboratory for her guidance on HPLC and Dr. Bhaskar Godogu from the mass spectrometry facility at the University of Pittsburgh for providing advice on HPLC-MS in this study. We thank the Alex Deiters lab at the University of Pittsburgh for sharing the Tecan 100 plate reader that supported measurements in this study. We thank Dylan T. Tomares of the Childers laboratory for his comments on the figures.

## AUTHOR CONTRIBUTIONS

W.S.C., C.Z., and J.W. conceptualized and designed the study. C.Z. and J.W. constructed plasmids and strains. C.Z. performed and collected preliminary data for I3A dose-response and ligands screening experiments on parent strains, pDawn-I3A, and pDusk-I3A_p1; J.W. performed the experiments and analyzed data. C. Z. performed and analyzed LC-MS experiments. W.S.C. and J.W. drafted the manuscript with input from C.Z. All authors provided critical feedback and helped shape the research, analysis, and manuscript.

## ASSOCIATED CONTENT

### Supporting Information

Figure S1-S4: Structure of *HtlD*^*I3A-PAS*^ and design of chimera I3A-HK, construction of I3A sensors, the selectivity of the I3A sensors, quantification of I3A and IND in the cell culture using LC-MS, and indole metabolite stimulation of homologs of HtlD^I3A-PAS^. Tables S1-S8: PDB PAS domains with ligands, promoters used for constructs, selected homologs of HtlD^I3A-PAS^, DNA oligos used in the study, Gibson cloning strategies for plasmids, *E*.*coli* strains used in the study, BLAST results for HtlD^I3A-PAS^, and chemicals used in the study.

